# Deep Learning Enhanced Fast Fluorescence Lifetime Imaging with A Few Photons

**DOI:** 10.1101/2023.04.06.534322

**Authors:** Dong Xiao, Natakorn Sapermsap, Yu Chen, David Day-Uei Li

## Abstract

We present a deep learning (DL) framework, which we term FPFLI (**F**ew-**P**hoton **F**luorescence **L**ifetime **I**maging), for fast analyzing fluorescence lifetime imaging (FLIM) data under highly low-light conditions with only a few photon-per-pixels (PPPs). FPFLI breaks the conventional pixel-wise lifetime analysis paradigm and fully exploits the spatial correlation and intensity information of fluorescence lifetime images to estimate lifetime images, pushing the photon budget to an unprecedented low level. The DL framework can be trained by synthetic FLIM data and easily adapted to various FLIM systems. FPFLI can effectively and robustly estimate FLIM images within seconds using synthetic and experimental data. The fast analysis of low-light FLIM images made possible by FPFLI will promise a broad range of potential applications.

## 1. Introduction

Fluorescence Lifetime IMaging (FLIM) is essential in diverse disciplines, including chemistry, pharmacy, fundamental biomedical science, and clinical applications [1]. FLIM uses the fluorescence lifetime of molecules to provide imaging contrast, allowing assessing cellular micro-environments such as pH, temperature, ion concentration, and metabolism [2, 3]. FLIM is also more robust than intensity imaging to measure Förster Resonance Energy Transfer (FRET), suitable for investigating protein-protein interactions and protein conformational changes [4, 5]. The time-correlated single photon counting (TCSPC) technique is the gold standard for FLIM measurement due to its high temporal resolution, high photon efficiency, and signal-to-noise ratio (SNR) approaching ideal Poisson statistics [6, 7]. A TCSPC generates decay histograms from time-stamped photons, from which fluorescence lifetimes are estimated.

However, obtaining enough photons to quantify fluorescence lifetimes precisely has been challenging. Traditional FLIM analysis is performed using least-square fitting (LSF) [8], maximum likelihood estimation (MLE) [9, 10], Bayesian analysis (BA) [11], Phasor methods [12, 13], or Centre-of-Mass Method (CMM) [14]. However, they can only produce reliable results with a sufficient photon count. For example, LSF requires at least a thousand photon-per-pixels (PPPs) for accurate multi-exponential decay analysis [9]. MLE and BA perform better for photon starvation scenarios with several hundred PPPs, but they suffer from a slow and complex analysis due to prolonged optimization routines [10, 11]. In addition, recent progress in deep learning (DL) approaches shows encouraging results for low-light FLIM with hundreds of PPPs [15-18]. DL offers a data-driven approach for fast analysis by directly mapping decay histograms to target lifetime parameters. However, as the detected PPP decreases to only several or dozens, it is mathematically impossible to recover the fluorescence lifetime using existing pixel-wise methods because the decay information within the pixel is permanently lost.

There are two reasons why obtaining sufficient photons can be challenging or even impossible in many scenarios. Firstly, the microenvironments can be photon-starved due to a low probe concentration, a poor quantum yield and photosensitivity in cells, and unoptimized excitation and emission wavelengths for FLIM measurements. Transporting or transferring fluorescent molecules can also result in low transporting/transfer efficiency for exogenous fluorophores, resulting in a low fluorescence intensity [19, 20]. Secondly, many FLIM measurements are strictly time-constrained. Excitation laser pulses can quickly reduce cell vitality and cellular reproduction [21] and lead to photobleaching and cell lysis [22]. If maintaining biological viability and avoiding photo-perturbation is necessary, the measurement time should be as short as possible with the laser power kept low.

Additionally, scientists are keen to observe rapid biomolecular interactions, requesting a much quicker acquisition. For example, characterizing fast-moving cell samples in high-throughput flow cytometers also poses a challenge due to the short dwell time [23]. Therefore, despite FLIM’s versatility, its application range is still limited by the required high photon budget.

To address these challenges, we propose a DL framework called Few-Photon Fluorescence Lifetime Imaging (FPFLI), breaking the conventional pixel-wise analysis paradigm to perform robust analysis in extremely low-light conditions. FPFLI recognizes the spatial correlation of fluorescence lifetimes among neighboring pixels and the information in fluorescence intensity images. We hypothesize that the information in the intensity image can benefit the lifetime estimation in a FLIM image. To tackle the severely ill-posed lifetime estimation problem caused by an ultralow PPP in a single pixel, FPFLI starts by estimating the average lifetime of the local area using a dedicated neural network. Then FPFLI uses another neural network to fuse the intensity and local lifetime information to reconstruct the fluorescence lifetimes for all pixels. We developed a large-scale synthetic FLIM dataset for fast training of the FPFLI algorithm. We demonstrate that FPFLI is rapid and robust for lifetime estimation that can push FLIM’s photon budget to a record low level with only a few PPPs.

## 2. Methods

### Algorithm implementation

FPFLI involves two main steps for estimating fluorescence lifetime images from low-light FLIM data, as illustrated in Fig. 1(a). Before lifetime estimation, we filter out noise background pixels and retain sample morphology through an image mask ***X***. In TCSPC measurement, the Poisson noise from the photon detection process plays a dominant role, and noisy background pixels can be easily distinguished using a lower threshold. Given the measured FLIM data ***H***(***i***, *t*), where ***i*** = [*i*_*x*_, *i*_*y*_] ∈ [1, *M*]^2^ is the spatial coordinate vector and *t* ∈ [1, *N*] is the temporal coordinate. The masked image is obtained by ***H***′ = ***X*** · ***H***. The dot operator denotes the scalar product of the matrices.

**Fig. 1.**
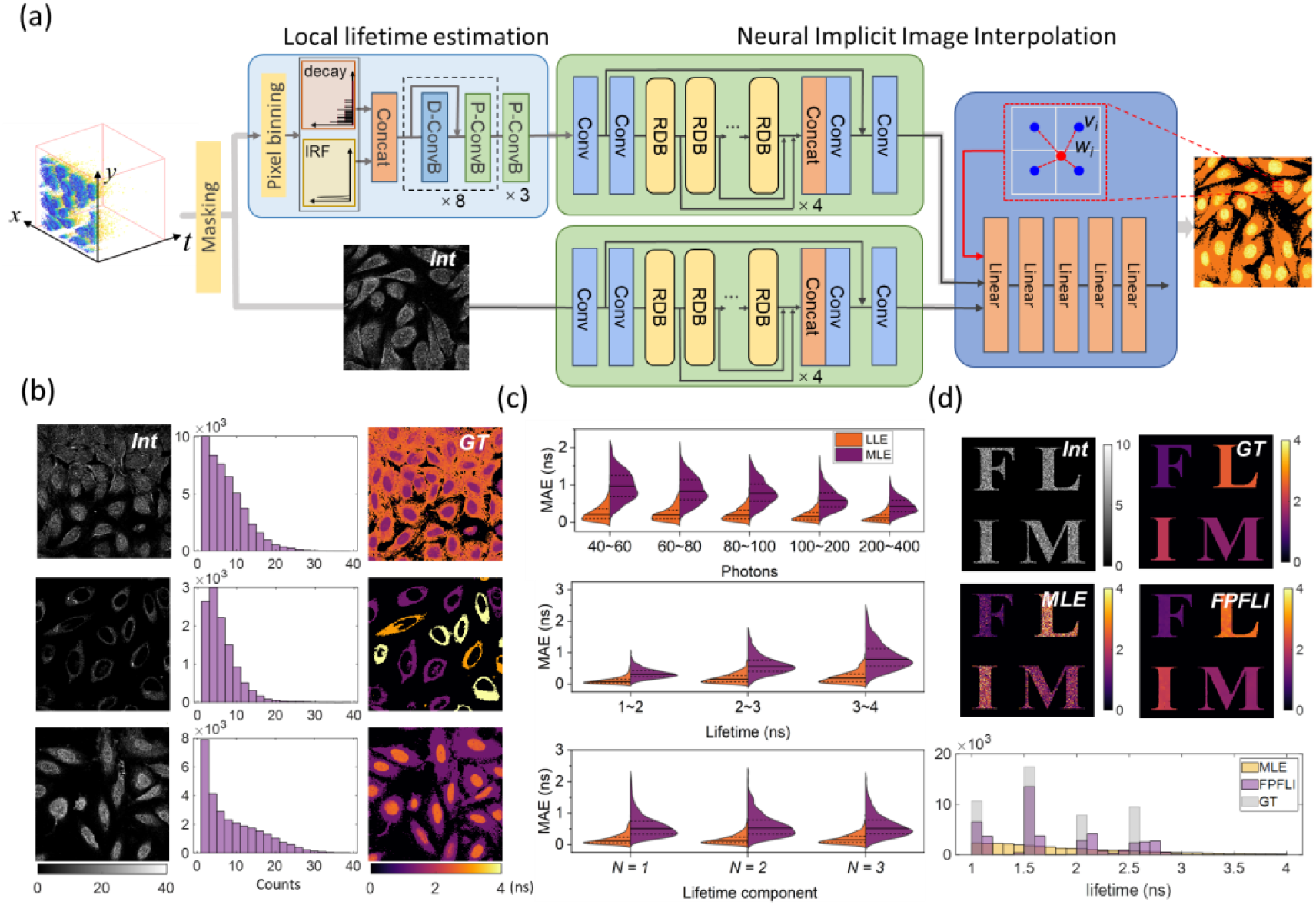
Algorithm overview of FPFLI. (a) FPFLI consists of two sub-networks: (i) Local Lifetime Estimator (LLE) using ConvMixer architecture for estimating the average lifetimes of local areas, and (ii) Neural Implicit Image Interpolation (NIII) network for reconstructing lifetime images. NIII contains an encoder and decoder. The encoder adopts a residual dense network (RDN) as the backbone, and the decoder is a five-layer dense network. (b) Semi-synthetic low light FLIM data for algorithm training. The average PPP of the samples range from 4 to 10, and the lifetime ranges from 1 to 4 ns. (c) Accuracy analysis of LLE under low light conditions. The ranges of photons, lifetime, and lifetime components are 40∼400, 1∼4 ns, and 1∼3, respectively. When one variable is under investigation, other variables change in their corresponding ranges randomly. The solid line indicates the median value, and the two dashed lines indicate the upper and lower quartiles. (d) NIII for analyzing synthetic low-light alphabetical FLIM images. The four letters have four lifetimes increasing from 1 to 2.5 ns with a 0.5 ns interval.

The first step of FPFLI is to estimate lifetime distributions in small local areas. As estimating the lifetime of individual pixels is no longer available, we utilize the strong spatial correlation of fluorescence lifetime images and perform pixel binning on neighboring pixels to evaluate the lifetime in that local area. For example, we merge the photon counts in an 8×8-pixel area into a single histogram with tens to hundreds of photons, which makes the local lifetime estimation well-posed. The FLIM data after spatial pixel binning is calculated as ***H***^*l*^ = 1^*k*×*k*×1^ ⊗ ***H***′, where *k* is the binning factor, ⊗_*s*=(*k,k*,1)_ is the three-dimensional cross-correlation operator with the stride *s* = (*k, k*, 1) along the spatial and temporal dimensions. To estimate the average lifetime in each local area, we use a Local Lifetime Estimator (LLE), which employs a one-dimensional (1D) ConvMixer architecture as its backbone [24] (**Suppl. note 1 and Fig. S1**). This lightweight architecture has fewer parameters, making it accurate and quick to train. To increase its applicability to different TCSPC FLIM systems, LLE takes both the instrument response function (IRF) and decay histograms as inputs. We denote the function of LLE as *C*(***θ***;·) with parameters ***θ***. The local lifetime image ***L***^***l***^ is calculated as:

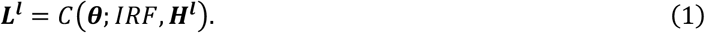

The second step is fusing the local lifetime distribution ***L***^***l***^ and the intensity image ***I*** to reconstruct the target lifetime image ***L***. The intensity image can be quickly obtained by summing up the photon counts in each pixel, as given by 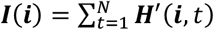. However, there is a size and pixel resolution mismatch between ***L***^***l***^ and ***I***. To address this, the neural implicit image interpolation (NIII) technique is used, which consists of an encoder and a decoder [25-28]. The encoder parameterizes the input images to latent codes using implicit neural representations (INRs). INRs use a continuous and differentiable implicit function to map coordinates to the corresponding signal, efficiently capturing signals’ underlying features with fewer parameters. Signals parameterized by INR are independent of spatial resolutions and can theoretically be sampled at arbitrary spatial resolutions. Encoder-based methods are proposed to share knowledge of all inputs instead of fitting individual functions for each input instance. In our study, two identical encoder networks are applied to extract two sets of latent codes distributed in spatial dimensions from ***L***^***l***^ and ***I***. The encoders adopt residual dense network (RDN) as the backbone to learn the local and global hierarchical features and extract latent codes [29] (**Suppl. note 1 and Fig. S1**). The sets of latent codes are expressed as:

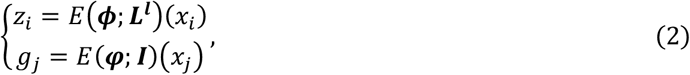

where *z*_*i*_ and *g*_*j*_ are the latent codes from ***L***^***l***^ and ***I*** for the query pixel coordinates *x*_*i*_ and *x*_*j*_, respectively. *E*(***Φ***; ·) and *E*(***φ***; ·) are the encoders with parameters ***Φ*** and ***φ***, respectively.

Once the latent codes are generated from the encoders, they are used as inputs along with query coordinates to a five-layer dense neural network decoder *F*(***σ***,) with parameters ***σ*** to generate weights and values for interpolating the target lifetime image ***L*** (**Suppl. note 1 and Fig. S1**). Given a query pixel coordinate *x*_*q*_ in ***L***, *a*_*q,i*_ and *v*_*q,i*_ are the interpolation weight and value between *i* and *q*, where *i* is a neighboring pixel for *q*. They are calculated as [28]:

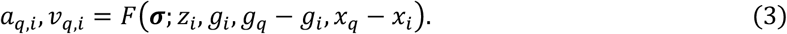

We chose the four nearest corner pixels of *q* in ***L***^***l***^, as shown in the insert in Fig. 1(a), as the set of neighbor pixels for *q*, denoted as **𝒩**_*q*_. The normalized weights *w*_*q,i*_ are obtained by applying the *SoftMax* function on *a*_*q,i*_:

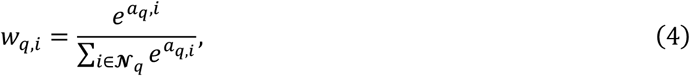

Then, the lifetime in *x*_*q*_ is calculated as:

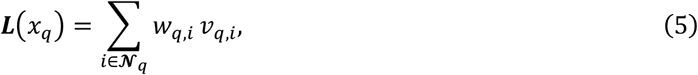

The reconstructed lifetime image ***L*** is obtained by iterating through all pixels.

### Semi-synthetic FLIM datasets

Developing DL algorithms for FLIM analysis faces a significant challenge due to insufficient FLIM datasets for training. Experimentally acquiring diverse FLIM images is time-consuming and labor-intensive, and the available samples often limit the cellular morphologies and decay features. Additionally, it is difficult to precisely control the SNR, lifetime component number, and lifetime ranges in experiments. Furthermore, FLIM images require a large photon count. Then they use a conventional algorithm to calculate the ground truth (GT) images, which further increases the experimental difficulty and limits the performance of DL algorithms.

We generated model-based semi-synthetic FLIM data to overcome these difficulties for algorithm training. We used immunofluorescence intensity images from the Human Protein Atlas (HPA) dataset [30]. We converted them into large-scale low-intensity FLIM datasets by considering both spatial Poisson distribution and temporal non-homogeneous Poisson process (**Suppl. note 2 and Fig. S2**). Without losing generality, we generated 5,000 low-light FLIM samples with a size of 256×256×256 for training purposes. The average PPP for pixels of the sample ranges from 4 to 10. The decay histograms in the training samples have a period of 10 ns and dynamic lifetime ranges from 1 to 4 ns. The training FLIM dataset covers diverse cellular morphologies, complex decay features, and varying lifetime distributions. Fig. 1(b) shows three low-count FLIM images from our training dataset. The first column displays the intensity maps of the FLIM images, whereas the second column presents the histogram of the intensity distribution. The intensity values range from zero to dozens, and most pixels only have a few photons, resulting in a low PPP of less than 10. The third column shows the ground truth (GT) lifetime maps, which are the amplitude-averaged lifetimes of all lifetime components.

### Network training and evaluation

We trained the LLE and NIII sub-networks separately. LLE was trained by synthetic decays with 256-time bins and a period of 10 ns. (**Suppl. Fig. S2**). The lifetime and photon count range from 1 to 4 ns and 50∼1000 counts, respectively. The lifetime components are randomly set to 1 to 3, with lifetime components randomly set between 1 to 3. To cover a wide range of TCSPC systems, we considered two types of IRFs, Gaussian and exponential, with FWHM ranging from 10 to 40 ps and peak position from the 10^th^ to 30^th^ time bin [31]. The training dataset comprised 100,000 samples. We used the amplitude-weighted average lifetime as the ground truth target, denoted as 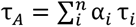, where *α*_*i*_ and τ_*i*_ represent the fraction ratio and lifetime of *i*^th^ lifetime component. This approach enabled us to include a broader range of cases without the concern of model mismatch problems that can arise when selecting specific exponential decay models, such as single- or bi-exponential models. Moreover, the amplitude-weighted average lifetime is robust for analyzing diverse applications [32]. We use the L_2_ norm as the loss function:

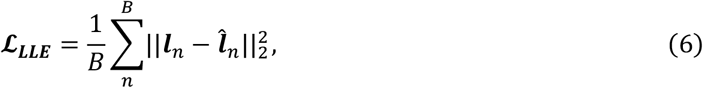

where *B* is the batch size, ***l*** and 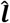 are the predicted and GT lifetimes, respectively. To train the NIII sub-network, we first processed the FLIM data using a trained LLE to estimate the local lifetime image ***L***^***l***^, which was then fed into the NIII with the intensity image ***I***. The lifetime deviation introduced by LLE was treated as a noise for the inputs to improve the training robustness. The spatial pixel binning was performed using 8×8 pixel patches. Next, we sampled pixels from the GT lifetime image and queried the decoder with the corresponding pixel coordinates to predict the pixel values. The L_1_ loss is applied for optimization:

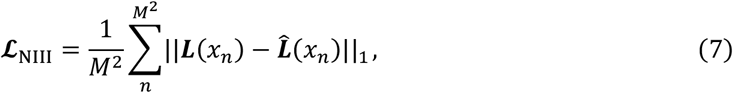

where *M*^2^ is the total pixel number in FLIM images. The detailed implementation and training are described in (**Suppl. note 1**). LLE required a significantly shorter training time, taking only 8 minutes, while NIII required a longer training time of approximately 60 hours for training.

After completing the training processes, we quantitatively evaluated the performance of both LLE and NIII. Figure 1(c) illustrates LLE’s performance on lifetime estimation under low-light conditions. Mean absolute error (MAE) was employed as a figure of merit to evaluate the accuracy of the algorithms, with the performance of MLE for lifetime estimation used as a benchmark for comparison purposes (The implementation of MLE is described in **Suppl. note 3**). We considered varying conditions, including different photon counts, lifetime ranges, and lifetime components. Results demonstrate that LLE significantly improved accuracy for lifetime estimation, with a much smaller median MAE and narrower distribution than MLE.

To evaluate NIII’s effectiveness and robustness, we also analyzed a 512×512 synthetic alphabetical FLIM image with an average PPP of 5. The image has distinct morphologic features, such as linear boundaries and sharp corners, that do not appear in training images. MLE failed to estimate the lifetime due to too fewer photons. In contrast, the reconstructed lifetime image by NIII is very close to the ground truth. The histogram further confirms the accuracy of lifetime estimation. The lifetimes estimated by FPFLI four have distinct clusters corresponding to the four GT lifetimes, while that for MLE have a flat distribution. It is worth noting that both LLE and NIII have low accuracy in estimating decays with long lifetimes, as seen in Fig. 1 (c) and (d). This is reasonable because, for a given photon count, the photons tend to distribute in more time bins for decay histograms with longer lifetimes, making it challenging to estimate lifetime. Nevertheless, Fig. 1 (c) and (d) demonstrate that FPFLI is a robust and effective method for analyzing low-light FLIM images.

## 3. Results

### Validating FPFLI on synthetic images

We began by validating FPFLI on synthetic FLIM images, which are ideal for quantitative analysis due to their controllable photon counts and known ground truth lifetime images. We generated a particular FLIM dataset for testing purposes using the same process as the training dataset and then employed the trained FPFLI algorithm to calculate the lifetime images. Fig. 2 illustrates four examples with varying shapes, granularities, and lifetime distributions. MLE takes more than an hour for lifetime calculation, but the images produced by MLE are erroneous and far from ground truth images. The lifetimes have a random distribution within an extensive range, making it difficult to obtain meaningful information.

**Fig. 2.**
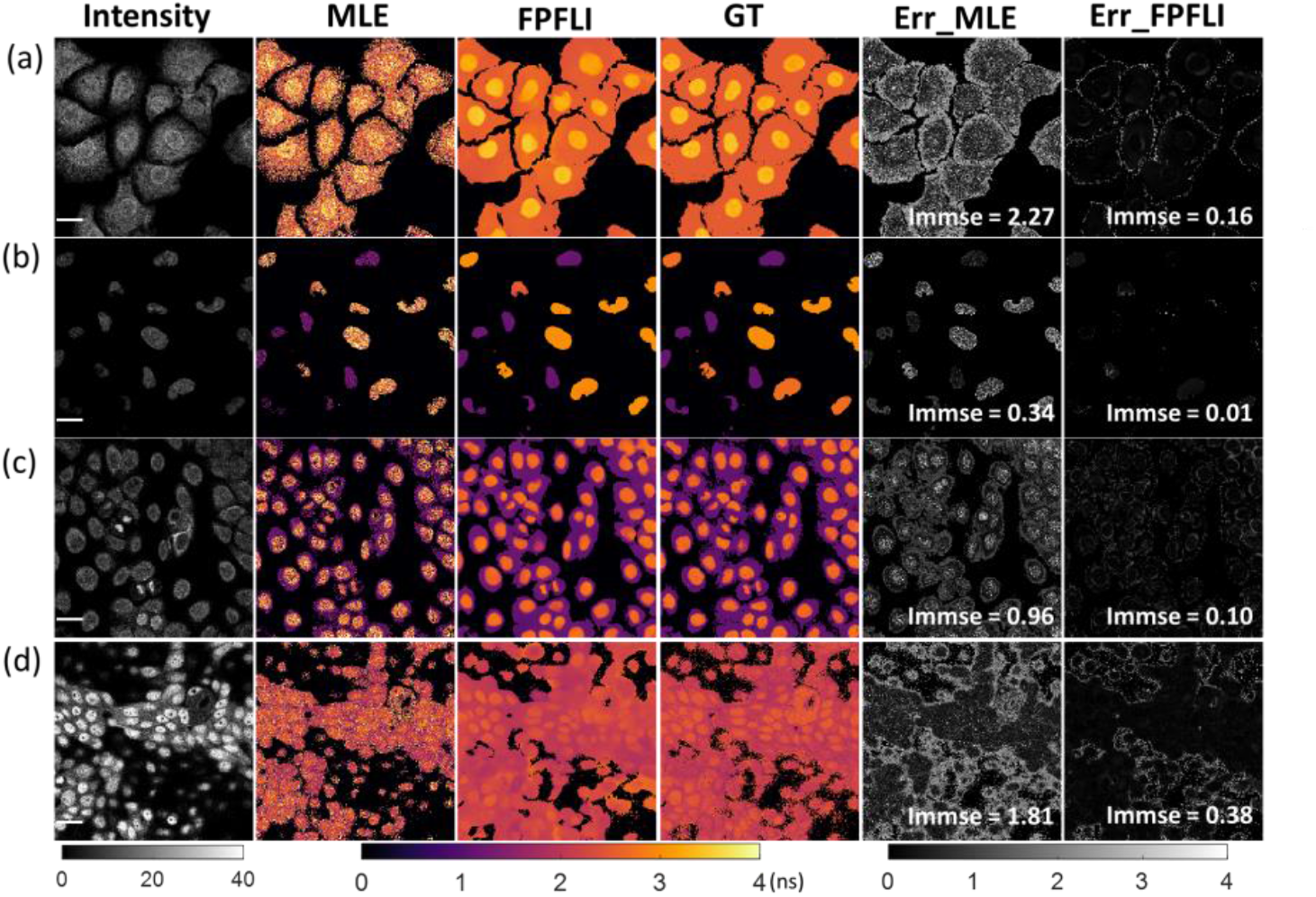
Validation of FPFLI on low-light synthetic FLIM images with data size of 256×256×256. (a) –(d) Four different samples and their analyzed results. The scale bar is 20 μm. The first column is the intensity images with the photon counts ranging from 0 to 40 (PPP varies from 5 to 10). In contrast, the second to fourth columns show the corresponding estimated lifetime images calculated from MLE and FPFLI and the ground truth (GT) images. The last two columns are the error map for MLE and FPFLI, respectively.

In contrast, FPFLI can estimate the fluorescence lifetime images within one second, producing highly accurate results almost identical to the ground truth images. It can reconstruct complex local fluorescence lifetime distributions from different cellular organelles, even if the information is vague and indistinguishable in the intensity images. For instance, FPFLI can reconstruct the lifetime and shapes of cell nuclei, even when their shapes cannot be identified by human eyes, as seen in Fig. 2 (a), (c), and (d). In practical FLIM measurements, the images often have uneven intensity distribution due to different fluorophores’ emission efficiencies and their varying concentrations. This raises the question of whether our method is robust to intensity perturbation and can extract correct intensity information for lifetime reconstruction. To verify this, Fig. 2(c) and (d) show the FLIM samples with non-uniform intensities. The sample in Fig. 2(c) has occasional bright spots, and the sample in Fig. 2(d) has a significant intensity variation. Despite these irregular intensity changes, the reconstructed lifetime images are unaffected, demonstrating FPFLI’s effectiveness and robustness.

Like any algorithm tackling ill-posed problems, FPFLI may produce inaccurate predictions. The error maps shown in the last two columns provide a means to estimate where errors are most likely to occur, defined as 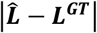 with 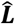 the predicted lifetime images by MLE or FPFLI. As expected, almost every pixel has a significant error for MLE, showing that MLE cannot converge to the correct lifetime under such low light conditions. In contrast, the error map for FPFLI has minor errors, mainly occurring at the boundaries. This is understandable because low-intensity images tend to shrink as the photons at boundaries are missing. We quantify the error using the image mean-squared error (IMMSE) as the figure of merit (*immse function*, MATLAB). A smaller IMMSE value indicates greater similarity between the reconstructed and GT images. In accordance with previous results, IMMSE for FPFLI is much smaller for all examples, showing that FPFLI is an effective algorithm for fluorescence lifetime estimation under low light conditions. We verified more samples with different shapes and lifetime distributions to confirm that FPFLI has a stable performance with high accuracy (**Suppl. Fig. S3**). Notably, FPFLI offers a fast calculation speed owing to its distinctive lifetime reconstruction approach. FPLFI requires estimating only a few lifetimes of local areas and inferring the entire lifetime image via interpolation. We performed a quantitative analysis of the calculation time of FPFLI on FLIM images of different sizes. The reconstruction of 256×256 and 512×512 lifetime images only took 0.63s and 0.72s, respectively. Even for large-scale 1024×1024 FLIM images, the calculation time was around 7s (**Suppl. Fig. S4**). Fast and accurately estimating lifetime can improve the discrimination of cellular structures under low-light scenarios, leading to potential applications such as in vivo and real-time FLIM-based surgical guidance for tumor excision or tissue delineation [33, 34]. This is particularly valuable in situations where the fluorescence signals have low SNR.

### FPFLI analysis of mouse kidney section

We then tested FPFLI on experimental FLIM data obtained from a cryostat section of a mouse kidney stained with Alexa Fluor 488 wheat germ agglutinin (F24630, FluoCells™ Prepared Slide, Thermo Fisher, UK). We used a confocal microscopic FLIM system (MicroTime 200D, Picoquant, Germany) with a 483 nm picosecond diode laser operating at 40 MHz to measure the cell images. Fluorescence emissions were collected by a 63× water immersion objective and recorded by a single photon avalanche diode (SPAD) detector with a 530/30 nm bandpass filter. To obtain low-intensity images, we terminated the measurement when the maximum photon count reached 40. We also measured corresponding high-intensity images with a maximum photon count of 4000 and fitted them by MLE as GT images. The measured FLIM data has 512×512 pixels, and their histograms have a 20 picosecond (ps) timing resolution with a period of 50 ns. We post-processed the images to 512×512×256 with a 0.1 ns timing resolution through temporal binning. Three cell images with different field-of-views (FOVs) are shown in Fig. 3, and their PPPs are around 6 to 7. The sample shapes and decay profiles show vast differences from the training data. We adapted FPFLI by retaining LLE using training data with a 0.1 ns timing resolution to analyse better the experimental data, which took only 8 minutes. The synthetic IRF was used because the SPAD detector has a wavelength-dependent response (**Suppl. note. S5**) [35].

**Fig. 3.**
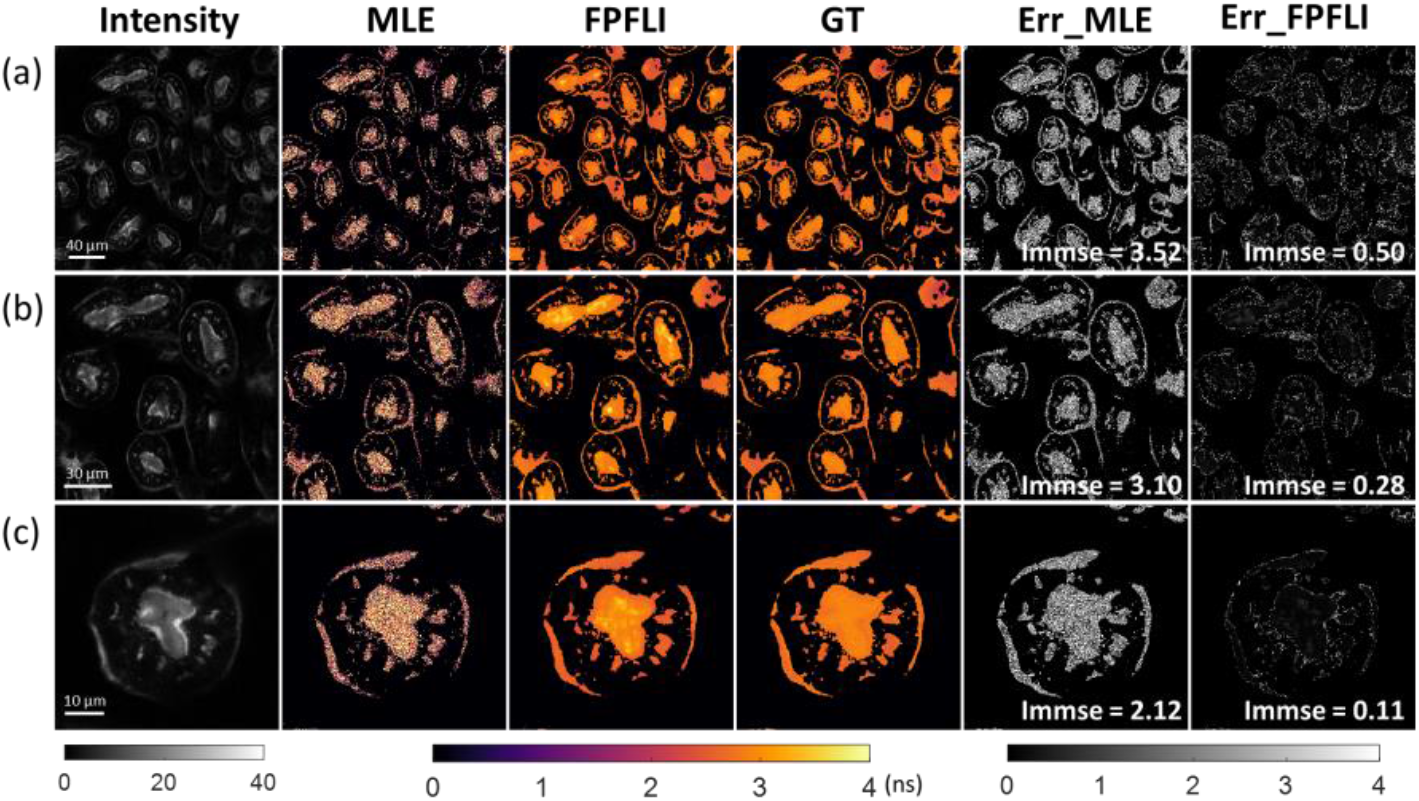
FPFLI analyzing FLIM images of the mouse kidney section with different FOVs. The average PPPs for (a) to (c) are 7.49, 6.78, and 7.46, respectively.

Consistent with previous findings, only FPFLI can produce meaningful lifetime images under the highly low-light conditions considered in this study. The overall lifetime distributions for all FOVs analyzed were in good agreement with their corresponding GTs, and subtle lifetime differences in some local areas were easily distinguishable. Quantitative analysis of IMMSE, similar to that for synthetic images, indicates that FPFLI trained by synthetic FLIM data has similar performance on different experimental FLIM data. The quick training of LLE enables FPFLI to adapt to varying decay histograms, making it a promising tool for general FLIM systems. However, it is worth noting that FPFLI can generate artefacts in some areas, such as the bright spots observed in the reconstructed lifetime images in Fig. 3(a)-(c). These artefacts are mainly due to relatively large lifetime deviations resulting from low photon counts, which can be eliminated by increasing the detected photons. In addition, images with high spatial complexity, such as those with complex shapes or lifetime distributions (Fig. 2(d) and Fig. 3(a)), have larger IMMSEs, indicating that FPFLI is more challenging to reconstruct lifetime images in such cases. Nevertheless, our results experimentally demonstrate that FPFLI can robustly estimate lifetime images from extreme low-light conditions with only a few PPPs, which is impossible for conventional FLIM analysis methods.

### Identifying the moving fluorophore-tagged microspheres

Analyzing fluorescence lifetime images under highly low light conditions allows imaging of fast-moving targets. To demonstrate this, we used FPFLI to analyze moving fluorophore-tagged microspheres, widely used in flow cytometry. Two kinds of fluorescent microsphere aqueous solution with a concentration of 3.6×10^6^ beads/ml, one containing Crimson (C, F8831, FluoSphere Polystyrene Microspheres, Thermo Fisher, UK) and the other one containing yellow-green (YG, F8836, FluoSpheres Polystyrene Microspheres, Thermo Fisher, UK) fluorescent microsphere, were mixed at an identical ratio. Both microspheres have an average size of 10 µm. The average lifetimes for C- and YG-microspheres, measured from their steady solution using a commercial lifetime fluorometer (FluoroCube Extreme, HORIBA Scientific, UK) were: *τ*_*C*_ from 1.9 to 2.1 and *τ*_*YG*_ from 2.9 to 3.2 ns, respectively. We dropped the mixed solution on a coverslip at room temperature to acquire the FLIM data and then scanned it using the MicroTime 200 FLIM system. The YG- and C-microspheres were excited using 40 MHz 483nm and 637nm picosecond pulsed lasers, respectively. The fluorescence light they emitted was detected by two separate SPAD detectors equipped with 530 nm and 650 nm long-pass filters, respectively. The microspheres underwent Brownian motion and slowly moved towards the boundary of the droplet due to the force of surface tension. To avoid motion blur, we kept the measurement time below 10 s. The measured FLIM data have 512×512 pixels and a 50 ns period with a 40 ps timing resolution. The data were then post-processed to 256 time bins with 0.1 ns timing resolution and analyzed by FPFLI.

Figs 4 (a) to (d) present the microsphere images with varying FOVs. Since it is difficult to accumulate high photon counts to calculate ground truth (GT) lifetimes for moving samples, GT images were obtained separately from different detector channels. The lifetimes of C- and YG-microspheres were assigned as *τ*_*C*_ = 2.0 and *τ*_*YG*_ = 3.0 ns for reference, respectively. Both microspheres have similar quantum yields (Fig. 4 (a)); therefore, it is impossible to identify the microsphere type from the intensity images. Additionally, as the photon counts are too few, lifetime images estimated by the traditional MLE also fail to recognise the microsphere type.

**Fig. 4.**
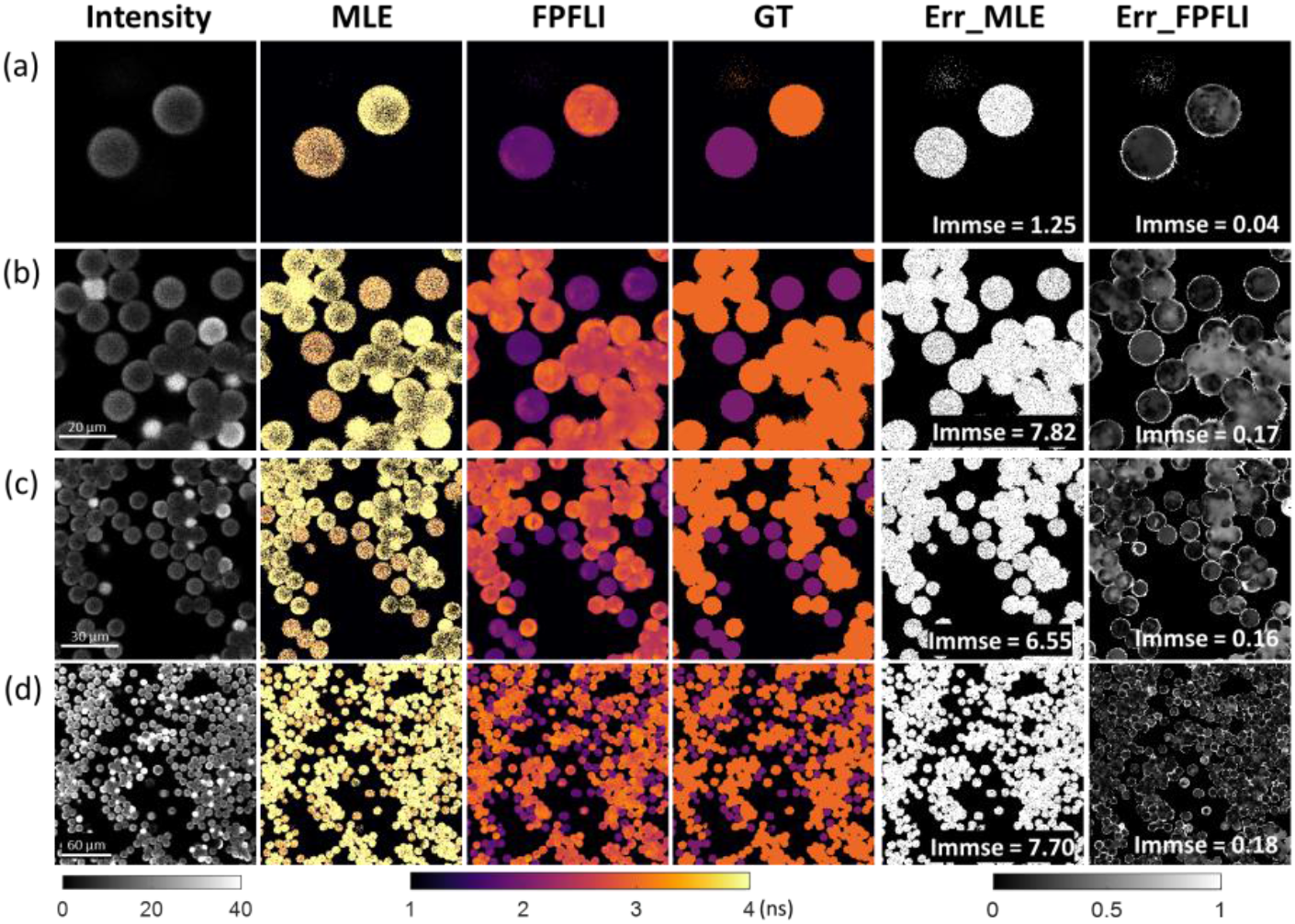
Identification of moving fluorophore-tagged microspheres with FPFLI. The average PPPs for microspheres in (a) to (d) are 8.4, 9.9, 9.2, and 15.7. The different quantum yields cause uneven intensity distributions in some microspheres, which were mixed from different production batches.

In contrast, lifetime images calculated by FPFLI quickly and accurately discriminate the microsphere types compared to the GT images. Even in the case of large FOV and extremely dense microsphere distribution (Fig. 4(d)), FPFLI maintains a high discrimination accuracy in identifying the types of microspheres. All C-microspheres can be sorted out from the lifetime image. The error maps and immse for FPFLI indicate that FPFLI has a small error for overall lifetime estimation. The bias for estimating the lifetime of C-microspheres is around ±0.1 ns. However, the estimated lifetime of YG-microspheres has an extensive distribution and skews towards larger values with a bias of 0.3 ns. Under the same photon count conditions, it is more challenging for FPFLI to estimate longer lifetimes, as discussed in the previous section (Fig. 1 (c)). To verify the robustness of FPFLI, we conducted a follow-up experiment using a two-photon FLIM system. In this setup, both microspheres were excited by a single laser, and a photomultiplier detector recorded their fluorescence signals. The measured FLIM images have much lower PPPs and significant background noise. Our findings demonstrate that FPFLI exhibits exceptional accuracy and robustness. Even for FLIM images with only one PPP, FPFLI still delivers lifetime images that can discriminate the types of microspheres (**Suppl. note 6 and Fig. S6**). This indicates that FPFLI holds excellent potential for high-throughput FLIM applications.

### FPFLI analyzing gold nanoparticle labelled Hek293 cells

The success of FPFLI in accurately estimating lifetimes depends on the spatial correlation of FLIM, which transforms the highly ill-posed problem into a well-posed one. As a result, DL neural networks make it possible to reconstruct lifetime images. However, some readers may question the performance of FPFLI when the spatial correlation in FLIM images is weak. To address this concern, we experimented using FPFLI to analyze the lifetime changes in Human Embryonic Kidney (Hek293) cells labeled with gold nanorods (GNRs). GNRs are promising contrast agents in biomedical research that offer several advantages over organic dyes [36-38]. These advantages include better photo-stability, low toxicity, extraordinary intracellular stability, broad-range fluorescence quenching capability, and the ability to conjugate to biomolecules. Their strong two-photon luminescence and characteristic short fluorescence lifetime are especially beneficial for FLIM imaging [39, 40]. Our experiments obtained the FLIM image of GNR-labeled Hek293 cells using a two-photon FLIM system. The LLE was quickly retrained for the system with a dynamic range from 1 to 3 ns. The sample preparation and image acquisition are described in (**Suppl. note 7**).

Fig. 5 (a) shows the low-light intensity image. Due to the strong two-photon luminescence, GNRs show bright spots with dozens of photons, irregularly scattered among cells. Nevertheless, most pixels of cells have fewer than 10 photons. In Fig. 5 (b), the corresponding GT lifetime image shows that GNRs shorten the cells’ lifetime because of their ultra-short lifetimes and fluorescence quenching effects. Due to the nanoscale size of GNRs, the lifetime changes occur at small discrete areas where GNRs accumulate and some independent single pixels. Therefore, the lifetime image has a low spatial correlation. Figs .5 (c) to (d) present the estimated lifetime images and their lifetime distributions obtained using MLE and FPFLI. Only FPFLI obtains a lifetime image with a lifetime distribution highly overlapped with that of the GT image. The low-lifetime regions identify the locations of GNRs. However, the image delivered by FPFLI becomes smooth and blurred compared to the GT image, resulting in more pixels with smaller lifetimes. This can be observed in both Figs. 5 (d) and (e).

**Fig. 5.**
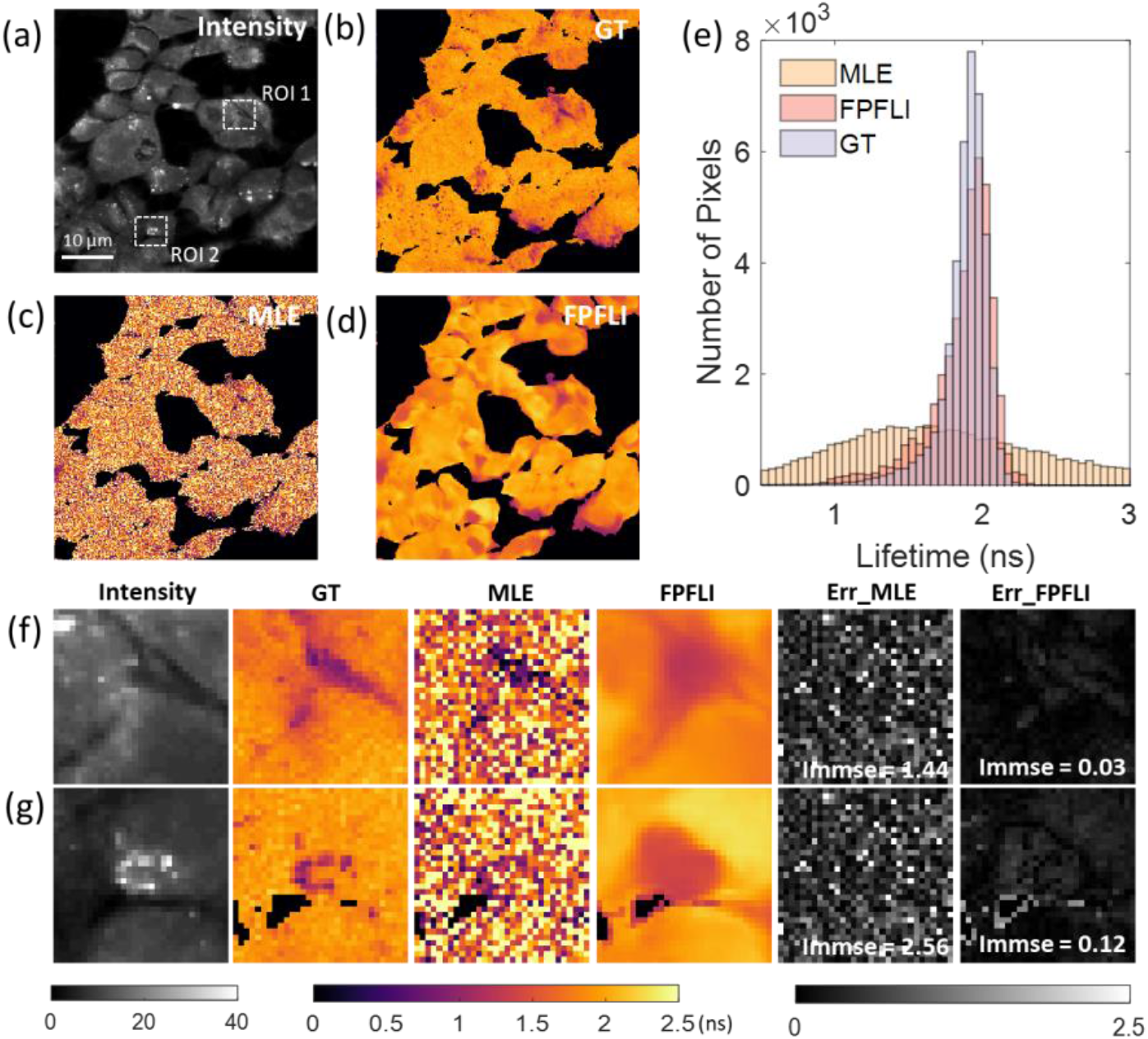
Fluorescence lifetime analysis of 256×256 gold nanorod-labelled Hek293 cells. (a) The intensity image of the cell sample. The measurement lasted for several seconds, and the average PPP of cells was around 9. White dashed boxes mark two ROIs with 31×31 pixels for detailed analysis. (b) The GT lifetime image was calculated from the same FLIM image with a maximum photon count of 10^4^. (c) and (d) The corresponding lifetime images for the low-light cell image calculated by MLE and FPFLI, respectively. (e) The lifetime distribution histograms for MLE, FPFLI, and GT. (f) and (g) Image comparisons of intensity, lifetime, and error maps in two ROIs.

For better analysis, Figs. 5 (f) and (g) provide detailed comparisons of intensity and lifetime images in two distinct regions of interest (ROIs). In ROI 1, the dark pixels show shorter lifetimes. In contrast, a bright spot at the left corner does not affect the lifetime, possibly due to GNRs being located inside the cells and the spot being caused by contaminations. On the other hand, the bright pixels corresponding to GNRs have shorter lifetimes in ROI 2. These two ROIs suggest that the high intensity is not always correlated to the short lifetime, and the pixels with shorter lifetimes are spatially discrete. Therefore, the intensity information cannot precisely guide the interpolation. FPFLI can only output lifetime images that are most likely to resemble the GTs, resulting in over-smoothed lifetime images. Despite that, the error maps of FPFLI indicate that the estimation accuracy remains high, with a deviation of about 0.1 and 0.3 for ROIs 1 and 2.

The results in Fig. 5 indicate that even in scenarios with low light and weak spatial correlation, FPFLI can still effectively estimate the overall lifetime distribution and local lifetime changes. However, it is essential to note that some artifacts, such as over-smoothed images and inaccurate lifetime estimations, may still occur. Therefore, it is crucial to be cautious to prevent incorrect interpretations of the results.

### 4. Discussion and Conclusion

We present FPFLI, a deep learning-based method to efficiently estimate fluorescence lifetime images under extreme low-light conditions. Our approach enables robust analysis for FLIM data with only a few PPPs without requiring slow and complex optimization processes. As a purely computational technique, FPFLI eliminates the need to modify FLIM systems or acquire additional data. The algorithm can be effectively trained using synthetic FLIM data and easily adapted for different experimental settings. We verified the effectiveness and robustness of FPFLI on both synthetic and experimental FLIM data obtained from confocal and two-photon FLIM systems.

FPFLI can be viewed as a DL-improved global analysis (GA) method. GA has been a popular choice for analyzing low-light FLIM data by accumulating all detected photons in a histogram for lifetime estimation and then assigning the same lifetime to all pixels of the sample [41]. GA provides a reliable lifetime readout with limited photons but sacrifices all the spatial information of fluorescence lifetime. A variant GA method incorporating image segmentation techniques can be used for analyzing spatially varying FLIM data [42]. However, this method is impractical due to the challenges in accurately segmenting images with low SNR, complex morphology, and uneven intensity distribution, as demonstrated in Figs. 2, 3, and 5. FPFLI addresses this difficulty by combining local lifetime estimation and global intensity image fusion. FPFLI’s sub-network LLE estimates the average lifetimes of small local areas like GA. Meanwhile, the information fusion with intensity image by NIII preserves the morphology information and provides both local- and non-local information to reconstruct the lifetime image. Thus, FPFLI retains the merit of GA while delivering as much spatially variant lifetime information as possible.

FPFLI employs a decoupled approach for local lifetime estimation and intensity image fusion, which brings significant advantages and flexibility. First, it reduces the algorithm complexity to a great extent. Unlike an end-to-end deep neural network that directly maps the 3D FLIM data to final lifetime images, FPFLI uses low-dimensional (1D and 2D) convolution operators, making the algorithm compact, easy to train, and faster. The unique lifetime reconstruction using NIII achieves unprecedented fast calculation speed, and the analysis of 512×512 images takes less than 2s. Second, FPFLI is more adaptable to different FLIM systems and application scenarios. For example, LLE can be retrained for decay histograms with varying timing resolutions and lifetime ranges in just a few minutes, as previously demonstrated (Fig. 3 and 4). In principle, FPFLI can be further adapted for time-gated FLIM systems, where the decay histograms are integrated into several time gates [43]. This allows for the extension of our algorithm to non-TCSPC systems.

Meanwhile, it is worth mentioning that NIII exhibits a robust performance for various images that show significant differences from the training data. This is due to the self-guided nature of NIII, as the intensity image is directly derived from the FLIM data. NIII can also be retrained using specified samples, enhancing its performance.

It is important to note that the size of the neighboring pixel for local lifetime estimation significantly impacts the algorithm’s performance. In this study, we utilized 8×8 patches to estimate local lifetimes for all considered FLIM data, which was found to be a good balance between the accuracy of local lifetime estimation and spatial complexity. If the local area under consideration is too small, for instance, 2×2 or 4×4 patches, the estimated local lifetimes tend to have low accuracy due to insufficient photon counts, which degrades the sequential intensity fusion. Conversely, suppose the local area is too large. In that case, the algorithm may fail to detect small local lifetime changes and output incorrect estimations and artefacts (**Suppl. Fig. S7**). Undoubtedly, no one-size-fits-all algorithm can consistently achieve excellent performance across all situations. The FPFLI framework can be flexibly modified and adjusted according to the specific properties of the FLIM data being analyzed. The LLE and NIII architectures could be replaced with more advanced alternatives.

In summary, our work presents FPFLI, a powerful tool for analyzing FLIM data with a few PPPs. FPFLI combines lifetime estimation of local areas and global information fusion with the intensity image, providing exceptional accuracy, fast speed, and high flexibility for adapting to different FLIM systems. FPFLI significantly reduces the photon budget with rapid analysis, promising broad and novel low-light FLIM applications previously unattainable with existing methods, such as investigating dynamic living samples and performing high-throughput imaging screens.

## Acknowledgements

This work was supported by Medical Research Scotland (1179-2017), the Biotechnology and Biological Sciences Research Council (BBSRC 20Alert SPRINT: BB/V019643/1), PicoQuant GmbH, and Photon Force Ltd.

## Notes

### Competing Interest Statement

The authors have declared no competing interest.

## Reference

1. J. R. Lakowicz, Principles of fluorescence Spectroscopy. (Springer, 2006).

2. K. Suhling, L. M. Hirvonen, J. A. Levitt, P. Chung, C. Tregidgo, A. L. Marois, D. A. Rusakov, K. Zheng, S. Ameer-Beg, S. Poland, S. Coelho, R. Henderson, and N. Krstajic, “Fluorescence lifetime imaging (FLIM): Basic concepts and some recent developments,” Med. Photon. 27, 3–40, (2015).

3. R. Datta, T. M. Heaster, J. T. Sharick, A. A. Gillette, and M. C. Skala, “Fluorescence lifetime imaging microscopy: fundamentals and advances in instrumentation, analysis, and applications,” J. Biomed. Opt. 25(7) 071203 (2020)

4. H. Wallrabe, and A. Periasamy, “Imaging protein molecules using FRET and FLIM microscopy,” Curr. Opin. Biotechnol. 16(1), 19–27 (2005).

5. Y. Sun, R. N. Day, and A. Periasamy, “Investigating protein-protein interactions in living cells using fluorescence lifetime imaging microscopy,” Nat. Protoc. 6, 1324 (2011).

6. W. Becker, Advanced Time-Correlated Single Photon Counting Applications. (Springer, 2015).

7. M. Wahl, “Time-Correlated Single Photon Counting”, Technical Note, PicoQuant GmbH (2014). https://www.picoquant.com/images/uploads/page/files/7253/technote_tcspc.pdf

8. M. Straume, S. G. Frasier-Cadoret, and M. L. Johnson, “Least-Squares Analysis of Fluorescence Data,”Topics in Fluorescence Spectroscopy, Chapter 4, pp. 177–240.Springer, 2002).

9. K. Santra, J. Zhan, X. Song, E. A. Smith, N. Vaswani, and J. W. Petrich, “What Is the Best Method to Fit Time-Resolved Data? A Comparison of the Residual Minimization and the Maximum Likelihood Techniques As Applied to Experimental Time-Correlated, Single-Photon Counting Data,” The Journal of Physical Chemistry B, 120(9), pp. 2484–2490 (2016).

10. T. A. Laurence and B. A. Chromy, “Efficient maximum likelihood estimator fitting of histograms,” Nat Methods, 7(5), pp. 338–389, (2010).

11. B. Kaye, P. J. Foster, T. Y. Yoo, and D. J. Needleman, “Developing and Testing a Bayesian Analysis of Fluorescence Lifetime Measurements,” PLoS One, 12(1), e0169337, (2017).

12. M. A. Digman, V. R. Caiolfa, M. Zamai, and E. Gratton, “The phasor approach to fluorescence lifetime imaging analysis,” Biophys J, 94(2) pp. L14–16, (2008).

13. C. Stringari, A. Cinquin, O. Cinquin, M. A. Digman, P. J. Donovan, and E. Gratton, “Phasor approach to fluorescence lifetime microscopy distinguishes different metabolic states of germ cells in a live tissue,” Proc. Natl. Acad. Sci. USA, 108(33), pp. 13582–13587, (2011).

14. D. D. Li, B. Rae, R. Andrews, J. Arlt, and R. K. Henderson “Hardware implementation algorithm and error analysis of high-speed fluorescence lifetime sensing systems using center-of-mass method,” J Biomed Opt 15(1), 017006 (2010).

15. J. T. Smith, R. Yao, N. Sinsuebphon, A. Rudkouskaya, N. Un, J. Mazurkiewicz, M. Barroso, P. Yan, and X. Intes, “Fast fit-free analysis of fluorescence lifetime imaging via deep learning,” Proc. Natl. Acad. Sci. USA 116(48), 24019–24030, (2019).

16. D. Xiao, Y. Chen, and D. D. Li, “One-Dimensional Deep Learning Architecture for Fast Fluorescence Lifetime Imaging,” IEEE J. Sel. Top. Quantum Electron 27(4), 1–10, (2021).

17. D. Xiao, Z. Zang, N. Sapermsap, Q. Wang, W. Xie, Y. Chen, and D. D. U. Li, “Dynamic fluorescence lifetime sensing with CMOS single-photon avalanche diode arrays and deep learning processors,” Biomed. Opt. Express 12, 3450–3462 (2021)

18. Y. I. Chen, Y. J. Chang, S. C. Liao, et al., “Generative adversarial network enables rapid and robust fluorescence lifetime image analysis in live cells,” Commun Biol 5, 18 (2022).

19. P. Kreiss, P. Mailhe, D. Scherman, B. Pitard, B. Cameron, R. Rangara, O. Aguerre-Charriol, M. Airiau, J. Crouzet, “Plasmid DNA size does not affect the physicochemical properties of lipoplexes but modulates gene transfer efficiency,” Nucleic Acids Research, 27(19), pp. 3792–3798 (1999).

20. P. Urayama, M. Mycek, “Fluorescence Lifetime Imaging Microscopy of Endogenous Biological Fluorescence,” Handbook of Biomedical Fluorescence. (CRC Press, 2003).

21. K. König, “Multiphoton microscopy in life sciences,” Journal of microscopy, 200(2) pp. 83–104 (2001).

22. K. R. Rau, A. Guerra Iii, A. Vogel, and V. Venugopalan, “Investigation of laser induced cell lysis using time-resolved imaging,” Appl. Phys. Lett., 84(15), pp. 2940–2942 (2004).

23. J. Nedbal, V. Visitkul, E. Ortiz-Zapater, G. Weitsman, P. Chana, D. R. Matthews, T. Ng, S. M. Ameer-Beg, “Time-domain microfluidic fluorescence lifetime flow cytometry for high-throughput Forster resonance energy transfer screening,” Cytometry A, 87(2), pp. 104–18 (2015).

24. A. Trockman, J. Zico Kolter, “Patches Are All You Need?” arXiv:2201.09792 (2022).

25. Y. Chen, S. Liu, and X. Wang, “Learning continuous image representation with local implicit image function.,” in Proceedings of the IEEE/CVF Conference on Computer Vision and Pattern Recognition, pp. 8628–8638 (2021).

26. K. Genova, F. Cole, A. Sud, A. Sarna, and T. Funkhouser, “ Local deep implicit functions for 3d shape,” in Proceedings of the IEEE/CVF Conference on Computer Vision and Pattern Recognition, pp. 4857–4866, (2020).

27. V. Sitzmann, J. N. P. Martel, A. W. Bergman, D. B. Lindell, and G. Wetzstein, “Implicit Neural Representations with Periodic Activation Functions,” arXiv:2006.09661, (2020).

28. J. Tang, X. Chen, and G. Zeng, “Joint Implicit Image Function for Guided Depth Super-Resolution,” presented at the Proceedings of the 29th ACM International Conference on Multimedia, (2021).

29. Y. Zhang, Y. Tian, Y. Kong, B. Zhong, and Y. Fu, “Residual Dense Network for Image Super-Resolution,” in Proceedings of the IEEE Conference on Computer Vision and Pattern Recognition (CVPR), 2472–2481 (2018).

30. The Human Protein Atlas, https://www.proteinatlas.org/

31. W. Becker, The bh TCSPC Handbook 9th edition (2021) https://www.becker-hickl.com/literature/documents/flim/the-bh-tcspc-handbook/

32. Li Y, Natakorn S, Chen Y, Safar M, Cunningham M, Tian J and Li DD-U, “Investigations on Average Fluorescence Lifetimes for Visualizing Multi-Exponential Decays,” Front. Phys. 8:576862, (2020).

33. A. Alfonso-Garcia, J. Bec, S. S. Weaver, B. Hartl, J. Unger, M. Bobinski, M. Lechpammer, F. Girgis, J. Boggan, L. Marcu, “Real-time augmented reality for delineation of surgical margins during neurosurgery using autofluorescence lifetime contrast,” J. Biophotonics 13, e201900108 (2019).

34. M. M. Lukina, L. E. Shimolina, N. M. Kiselev, V. E. Zagainov1, D. V. Komarov, E. V. Zagaynova and M. V. Shirmanova, “Interrogation of tumor metabolism in tissue samples ex vivo using fluorescence lifetime imaging of NAD(P)H, “ Methods Appl. Fluoresc. 8(1), 014002 (2019).

35. D. Xiao, N. Sapermsap, M. Safar, M.R. Cunningham, Y. Chen, and D. D-U. Li, “On Synthetic Instrument Response Functions of TimeCorrelated Single-Photon Counting Based Fluorescence Lifetime Imaging Analysis,” Front. Phys. 9:635645 (2021).

36. E. Boisselier and D. Astruc, “Gold nanoparticles in nanomedicine: preparations, imaging, diagnostics, therapies and toxicity,” Chem. Soc. Rev., 38(6), pp. 1759–1782 (2009)

37. G. Wei, J. Yu, J. Wang, P. Gu, D.J.S. Birch, Y. Chen, “Hairpin DNA-functionalized gold nanorods for mRNA detection in homogenous solution,” J. Biomed. Opt. 21, 097001 (2016).

38. Y Chen, Y Zhang, DJS Birch, AS Barnard, “Creation and luminescence of size-selected gold nanorods,” Nanoscale, 4 (16), 5017–5022 (2012).

39. Y. Zhang, J. Yu, D.J.S. Birch, Y. Chen, “Gold nanorods for fluorescence lifetime imaging in biology,” Journal of biomedical optics 15 (2), 020504–020504-3 (2010)

40. Y Zhang, D.J.S. Birch, Y. Chen, “Two-photon excited surface plasmon enhanced energy transfer between DAPI and gold nanoparticles: opportunities in intra-cellular imaging and sensing,” Applied Physics Letters 99 (10), 103701 (2011).

41. P. J. Verveer, A. Squire, P. I.H. Bastiaens, “Global Analysis of Fluorescence Lifetime Imaging Microscopy Data,” Biophysical Journal, 78(4), pp. 2127–2137 (2000).

42. S. Pelet, M. Previte, L. Laiho, and P. So, “A fast global fitting algorithm for fluorescence lifetime imaging microscopy based on image segmentation,” Biophysical J. 87(4), pp. 2807–2817 (2004).

43. M.J. Cole, J. Siegel, S.E. Webb, R. Jones, K. Dowling, M. J. Dayel, D. Parsons-Karavassilis, P. M. French, M. J. Lever, L. O. Sucharov, M. A. Neil, R. Juskaitis, T. Wilson. “Time-domain whole-field fluorescence lifetime imaging with optical sectioning,” J Microsc. 203(3), pp. 246–57 (2001).

